# Diversity-dependent plant-soil feedbacks underlie long-term plant diversity effects on primary productivity

**DOI:** 10.1101/376269

**Authors:** Nathaly R. Guerrero-Ramírez, Peter B. Reich, Cameron Wagg, Marcel Ciobanu, Nico Eisenhauer

**Author notes:** **Statement of authorship:** N.E. conceived the idea, N.G-R and N.E developed the idea, P.B contributed with the first experimental phase, N.G-R implemented the study and collected data with the help of M.C. N. G-R analyzed the data and wrote the paper with substantial input from all authors. **Corresponding author:** Nathaly R. Guerrero-Ramírez.

## Abstract

Although diversity-dependent plant-soil feedbacks (PSFs) may contribute significantly to plant diversity effects on ecosystem functioning, the influence of underlying abiotic and biotic mechanistic pathways have been little explored to date. Here, we assessed such pathways with a PSF experiment using soil conditioned for ≥12 years from two grassland biodiversity experiments. Model plant communities differing in diversity were grown in soils conditioned by plant communities with either low- or high-diversity (soil history). Our results reveal that plant diversity can modify plant productivity through both diversity-mediated plant-plant and plant-soil interactions, with the main driver (current plant diversity or soil history) differing with experimental context. The underlying mechanisms of PSFs were explained to a significant extent by both abiotic and biotic pathways (specifically, nematode richness and soil nitrogen availability). Thus, effects of plant diversity loss on ecosystem functioning may persist or even increase over time because of biotic and abiotic soil legacy effects.

## Introduction

Increasingly positive plant diversity effects on ecosystem functioning over time have been found in experimental ecosystems (Cardinale *et al*. 2007; Reich *et al*. 2012; Guerrero-Ramírez *et al*. 2017). Temporal strengthening of the biodiversity-ecosystem functioning (B-EF) relationship can be driven, depending on the experimental context, by an increase in primary productivity in high-diversity communities, a decrease in primary productivity in low-diversity communities, or both (Guerrero-Ramírez *et al*. 2017). Therefore, temporal changes in the B-EF relationship depend not only on mechanisms underlying an increase of ecosystem functioning in communities with high-diversity, which has been the main focus in previous work (Cardinale *et al*. 2007; Reich *et al*. 2012), but also mechanisms underlying a decrease of plant primary productivity in communities with low-diversity (Meyer *et al*. 2016; Guerrero-Ramírez *et al*. 2017).

From a mechanistic perspective, a strengthening of the relationship between plant diversity and primary productivity may result from dynamic temporal interactions between plants and their soil biotic and abiotic environment (Eisenhauer 2012; van der Putten *et al*. 2013). High- and low-diversity plant communities may increase or decrease their productivity over time, respectively, by modifying positively or negatively their soil environment, i.e. by inducing positive or negative plant-soil feedbacks (PSFs). Positive PSFs may be the result of high-diversity plant communities allowing for the diversification of soil microhabitats that in turn likely promotes the diversity of soil organisms (Hooper *et al*. 2000; Eisenhauer *et al*. 2017). Highly diverse plant communities can also accumulate and exploit more soil nutrients than less diverse communities, creating a resource gradient with increasing plant diversity over time (Fargione *et al*. 2007; Oelmann *et al*. 2010, 2011). These changes in the soil biotic and abiotic environment may, in turn, feedback on primary plant productivity (Bever *et al*. 2010). For example, highly diverse soil communities may increase primary productivity by influencing resource partitioning *via* increasing the available biotope space (Dimitrakopoulos & Schmid 2004), and the available forms of soil nutrients (Turner 2008; Eisenhauer 2012). Alternatively, negative PSFs may be the result of plant communities with low-diversity accumulating soil communities dominated by plant antagonists (Maron *et al*. 2011; Schnitzer *et al*. 2011). Low-diversity plant communities may also use soil nutrients in an imbalanced way and thus deplete soil nutrient stocks (Schenk 2006). Thus, soil conditioned by low-diversity plant communities may not only have a negative effect on plant productivity in the long run, but also, when used as a baseline, may co-determine the B-EF relationship.

Evidence for the role of PSFs on the B-EF relationship has usually been compartmentalized by focusing on either the effect of plants on soil abiotic *or* biotic conditions (e.g. De Deyn *et al*. 2004; Fargione *et al*. 2007), or on the effect of soil abiotic *or* biotic conditions on ecosystem functioning (e.g. Fridley 2002; Wagg *et al*. 2014). Yet few studies have included the effect of plant communities on ecosystem functioning *via* soil, i.e. sequential plant-soil-plant responses. One of the few studies that included sequential responses highlighted a negative PSF in plant communities with low-diversity (Yang *et al*. 2015). These negative feedbacks were attributed to the accumulation of plant pathogens and herbivores in low-diversity plant communities. However, a direct approach identifying which mechanisms are underlying PSFs, such as specific changes in the soil biotic and abiotic environment are assessed directly rather than inferred indirectly, e.g. by sterilizing soils, is still missing.

The effect of plant communities on plant primary productivity *via* soil typically has been determined using responses at the species-level and, to a lesser extent, responses at the community-level. Evidence suggests that the direction and magnitude of PSFs may differ between levels of organization, with a stronger negative PSF at the species level as compared to the community level (Kulmatiski *et al*. 2008). However, caution is required due to the lack of available information at the community level. In the context of B-EF relationships, it is likely that a positive PSF may influence responses at the community level and not at the species level. Specifically, potential PSFs of soil conditioned by high-diversity plant communities, e.g. *via* increases in soil biodiversity and available nutrient forms, may be exploited more completely by diverse plant communities, e.g. through plant resource use complementarity (e.g. Loreau & Hector 2001; David Tilman *et al*. 2014), and not by low-diversity communities.

The impact of plant communities with low-diversity on ecosystem functioning *via* soil may differ based on the functional relatedness between the plant community that conditioned, and the plant community currently growing in the soil (Bezemer *et al*. 2006; Cortois *et al*. 2016). Sharing plant functional identity may amplify negative PSFs due to accumulation of specialized plant antagonist and/or depleting specific nutrient stocks. Alternately, sharing plant functional identity may amplify positive PSFs due to accumulation of specialized plant facilitators (Cortois *et al*. 2016). Therefore, not only plant diversity but also plant functional identity may play a key role in PSFs.

Nematode communities potentially provide a mechanistic link in PSFs (Cortois *et al*. 2017). Nematodes are commonly used as indicators of soil health and plant growth (Neher 2010). Nematode communities not only respond to differences in plant communities (De Deyn *et al*. 2004; Eisenhauer *et al*. 2011) but also provide information about potential functional roles and soil feedback effects on plant productivity, e.g. plant antagonists *versus* plant growth facilitators (Neher 2010; Cortois *et al*. 2017). Plant-feeding nematodes are important plant antagonists that are known to be able to cause substantial losses in crop yields (Neher 2010). Alternately, microbial-feeders, predators and omnivores are regarded as plant growth facilitators as they accelerate nutrient cycling and consumption of plant-feeders (Neher 2010). While microbial-feeders may suppress nutrient availability by affecting the abundance of microbes, the positive effects on nutrient cycling generally surpass the losses by stimulating microbial activity, recycling immobilized nutrients, and enhancing bioturbation (Ettema 1998).

Here, we aimed to assess the role of PSFs in the B-EF relationships in two long-term plant diversity experiments. To do so, we determined sequential plant-soil-plant responses using soil conditioned for more than ten years by plant communities with either low- or high-diversity (soil history). To consider multiple potential mechanistic pathways that underlie the effects of PSFs on plant productivity, we explored the roles of soil biotic (soil nematode communities) and abiotic (available soil nutrients) conditions. We hypothesized that plant species richness may influence primary productivity *via* dissimilar biotic and abiotic soil conditions. On the one hand, we expected positive effects of soil conditioned by high-diversity plant communities on plant productivity by increasing soil nematode richness, the density of plant-growth facilitating nematodes, and available soil nutrients. On the other hand, we expected a negative effect of soil conditioned by low-diversity plant communities through an increase in the abundance of plant-feeding nematodes and a reduced availability of nutrients. Moreover, we expected that the positive PSF of soil conditioned by high-diversity plant communities to be higher in plant species mixtures in the feedback phase than in plant communities with only one plant functional group. Finally, we expected the negative PSF of soil conditioned by low-diversity plant communities to be higher in low-diversity plant communities in the feedback phase growing in soil conditioned by their own plant functional group than in soil conditioned by other plant functional groups.

## Materials and methods

### Conditioning phase

Soil from two long-term grassland biodiversity experiments, the BioCON Experiment (Reich *et al*. 2001) and the Jena Experiment (Roscher *et al*. 2004), was used to establish a PSF experiment. The BioCON Experiment is located at the Cedar Creek Ecosystem Science Reserve in Minnesota, USA on sandy soil (Nymore series, Typic Upidsamment). The BioCON Experiment was planted in 1997 using a pool of 16 species including grasses, forbs, and legumes (Reich *et al*. 2001). The Jena Experiment is located in Jena, Germany on the floodplain of the River Saale, and the soil is Eutric Fluvisol. The Jena Experiment was sown in 2002 using a pool of 60 species including grasses, forbs, and legumes (Roscher *et al*. 2004).

From each biodiversity experiment, soil samples from 12 plots with one plant species (low-diversity plots) and 5 plots with either nine (BioCON Experiment) or eight (Jena Experiment) plant species (high-diversity plots) were collected in summer of 2014. Soil was conditioned for 17 and 12 years in the BioCON and Jena Experiments, respectively. Notably, in the BioCON Experiment, we only sampled plots with ambient atmospheric CO2 concentrations and N availability. For low-diversity plots, a minimum of three plots was selected for each plant functional group: grasses, forbs, and legumes. In each plot, three soil cores were taken (depth 20 cm, diameter 3.8 or 5 cm) and mixed gently in a plastic bag (between 600 and 800 g of fresh soil were collected per plot). Afterwards, the soil was sieved using a 4 mm mesh. From each soil sample, a sub-sample of 25 g of fresh soil was taken to extract nematodes using a modified Baermann method (Ruess 1995, Appendix S1 in Supporting Information). Nematodes were then grouped into different feeding groups: abundance of plant-feeders, bacteria-feeders, fungal-feeders, omnivores, and predators. This information was used in a principal component analysis (PCA) to determine variation in the functional composition of nematode communities among experimental plots (Figs. S1 and S2). PC axis two was used in the statistical analyses, because it represents a gradient from plant antagonists (plant-feeders) to plant growth facilitators (fungal-feeders) in both biodiversity experiments.

Plant available nitrogen (N) and phosphorus (P) were also quantified using sub-samples taken from the soil in the two biodiversity experiments. A proxy of extractable or available N (mg kg^-1^) was calculated by adding the N concentrations of ammonium (NH4) and nitrate (NO3; Appendix S1).

### Feedback phase

Soil collected from each plot was divided into four sub-samples to establish four new plant communities in a microcosm experiment: one plant community for each plant functional group (grasses, forbs, and legumes) and one plant community that contained all three plant functional groups (Fig. 1a). The feedback experiment was established in microcosms (diameter of 6 cm at the bottom and 9 cm at the top, and a height of 7 cm, Appendix S2). Two plant species for each plant functional group were selected for each experimental site. To avoid species-specific PSFs, the selected plant species used in the feedback experiment were present in the study area but were not present in the sampled plots (Appendix S2). Each microcosm contained six plant individuals to keep plant density constant among treatments. After transplanting, the experiment was run for six weeks.

**Figure 1.**
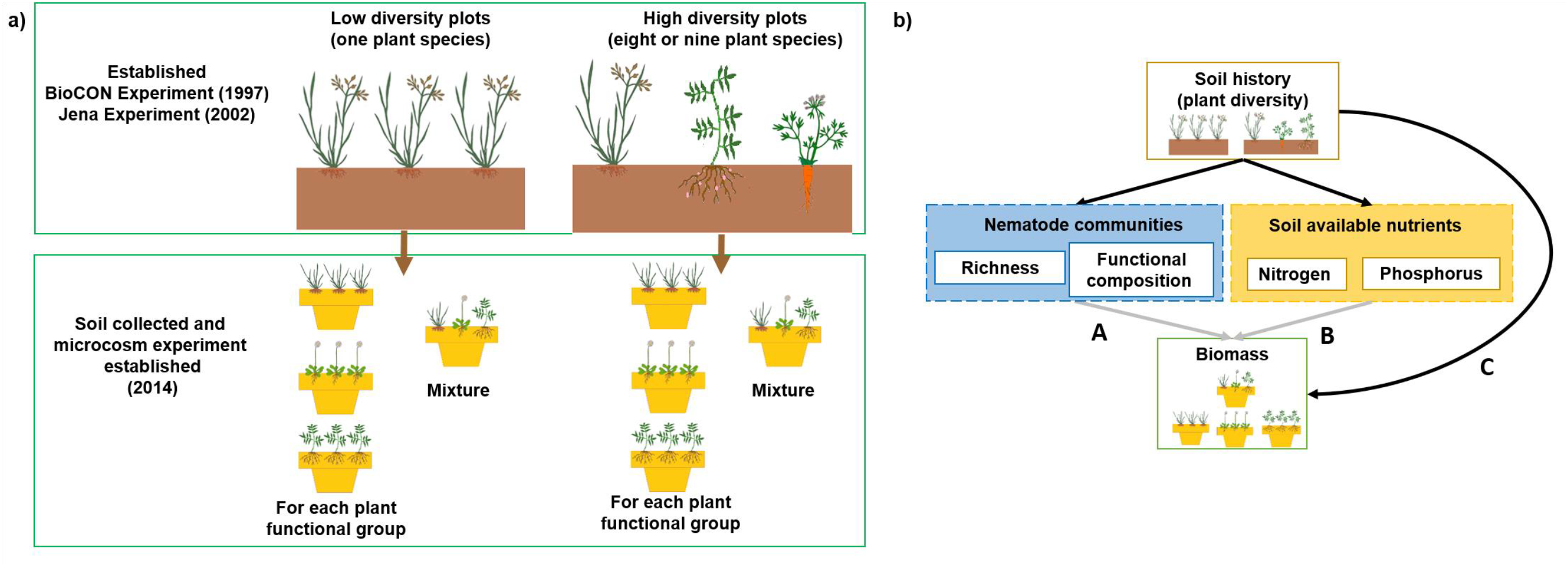
Experimental design of the present study **(a)**. Soils from two long-term biodiversity experiments, the BioCON and the Jena Experiments, were collected in 2014. Specifically, soil conditioned (for more than ten years) by plant communities with either low species diversity (12 plots with one plant species) or high species diversity (five plots either with eight or nine plant species for the Jena and the BioCON Experiments, respectively). Collected soils were used to establish four new plant communities, i.e. grasses only, forbs only, legumes only, and plant species mixtures of all three plant functional groups in a microcosm experiment. The plant-soil feedback experiment ran for six weeks. Hypothetical causal model explaining soil history effects **(b)**. Positive and negative plant-soil feedbacks on plant biomass may be explained by the long-term influence of plant diversity on biotic drivers, i.e. nematodes communities (A) and/or abiotic drivers, i.e. soil available nutrients (B). Mechanisms that were not captured by neither nematode community nor soil available nutrient pathways are represented by direct effects of soil history diversity on plant biomass (C).

In total, 136 microcosms were established: two studies (the BioCON and Jena Experiments) × 17 plots (12 soils conditioned by low- and five soils conditioned by high-diversity plant communities, i.e. soil history) × 4 plant communities (grasses only, forbs only, legumes only, and a mixture of all three plant functional groups). At the end of the experiment, shoot and root biomass was harvested and dried at 70°C for 72 h and weighed (0.001 g).

To determine plant diversity effects, proportional deviance was calculated (Loreau 1998, Appendix S3). To test whether non-additive diversity effects were found in plant mixtures, 95% confidence intervals (CI) were calculated using 10,000 bootstrap replicates (adjusted percentile bootstrap). Non-additive effects are considered to occur when the 95% CI do not overlap with zero.

Based on Baxendale *et al*. (2014), a PSF index was calculated to determine the role of plant functional identity of low-diversity communities (Appendix S4). To test whether plant functional identity affected PSFs, 95% CI were calculated using 10,000 bootstrap replicates (adjusted percentile bootstrap). Positive values indicated higher plant biomass in plant communities growing in soil conditioned by their own plant functional group than in soil conditioned by other plant functional groups. In contrast, negative values indicated higher plant biomass in plant communities growing in soil conditioned by other plant functional groups than in soil conditioned by their own plant functional group. Strong (positive or negative) PSFs are considered to occur when the 95% CI do not overlap with zero.

### Statistical analysis

The effects of soil history (conditioning phase), plant diversity in microcosms (feedback phase), and their interaction on total, shoot, and root biomass of plant communities in the microcosm experiment were determined with a linear mixed-effect model using the “nlme” package (Pinheiro *et al*. 2018). To account for potential heterogeneity in soil conditions (Reich *et al*. 2001; Roscher *et al*. 2004), plot nested in experimental blocks/rings were included as a random effect. To fulfill the assumptions of linear mixed-effect models, square-root transformations were used in the BioCON Experiment (Appendix S5).

To assess the influence of soil history on plant biomass operating *via* soil biotic and abiotic conditions, structural equation modeling (SEM) based on piecewise fitting of linear mixed-effect models using the “piecewiseSEM” package were performed (Lefcheck 2016). The SEM allowed us to test a hypothetical causal model based on *a priori* knowledge of PSFs (Fig. 1b). A direct path was included between soil history (plant diversity levels in the conditioning phase) and nematode communities, i.e. nematodes richness and functional composition (PC axis 2), and between soil history and soil available nutrients (N and P). As nematode communities and soil available nutrients are potential mechanistic pathways explaining PSFs, alternative paths between them and plant biomass were added, if this improved the model fit (based on modification indices, p-value < 0.05). Effects of soil history through mechanistic pathways were calculated by multiplying the effect of soil history on the biotic/abiotic explanatory variable and the effect of the biotic/abiotic explanatory variable on plant biomass in the microcosms. Mechanisms which were not capture by neither mechanistic pathways are represented by the direct path between soil history and plant biomass of plant communities in the microcosm experiment. Experimental blocks/rings were also included as a random effect. Independent path models were fitted for each biodiversity experiment (the BioCON and Jena Experiments). To fulfill the assumptions of the models in the BioCON and Jena Experiments, natural logarithmic and square-root transformations were carried out (Appendix S5). Model fits were assessed using Fisher’s C statistic based on the test of directed separation; if the test p-value was > 0.05, the data fits the hypothesized causal network (Lefcheck 2016). All analyses were performed using R 3.4.1 (RCoreTeam 2017).

## Results

### Effects of soil history, plant diversity, and their interaction on plant biomass

Soil history (conditioning phase) and the interaction between soil history and plant diversity in the microcosm experiment (feedback phase) influenced plant biomass in the BioCON Experiment (Table S1, Fig. 2). Plant biomass was higher in plant communities that grew in soil conditioned by more diverse plant communities (F_1,13_ = 6.059 and 6.316 for total and shoot biomass, respectively, p-values < 0.05; F_1,13_ = 3.450 for root biomass, p-value < 0.1). The positive effect of soil history on plant biomass in microcosms was stronger in plant mixtures than in plant communities with a single plant functional group (soil history × plant diversity in microcosms: F_1,48_ = 3.179 and 3.367 for total and shoot biomass, respectively, p-values < 0.1). In the Jena Experiment, soil history did not influence plant biomass in the microcosm experiment (Table S1). Instead, plant biomass increased with plant diversity in microcosms (F_1,49_ = 4.235 and 5.175 for total and root biomass, respectively, p-values < 0.05; F_1,49_ = 4.235 for shoot biomass, p-value < 0.1; Fig. 2).

**Figure 2.**
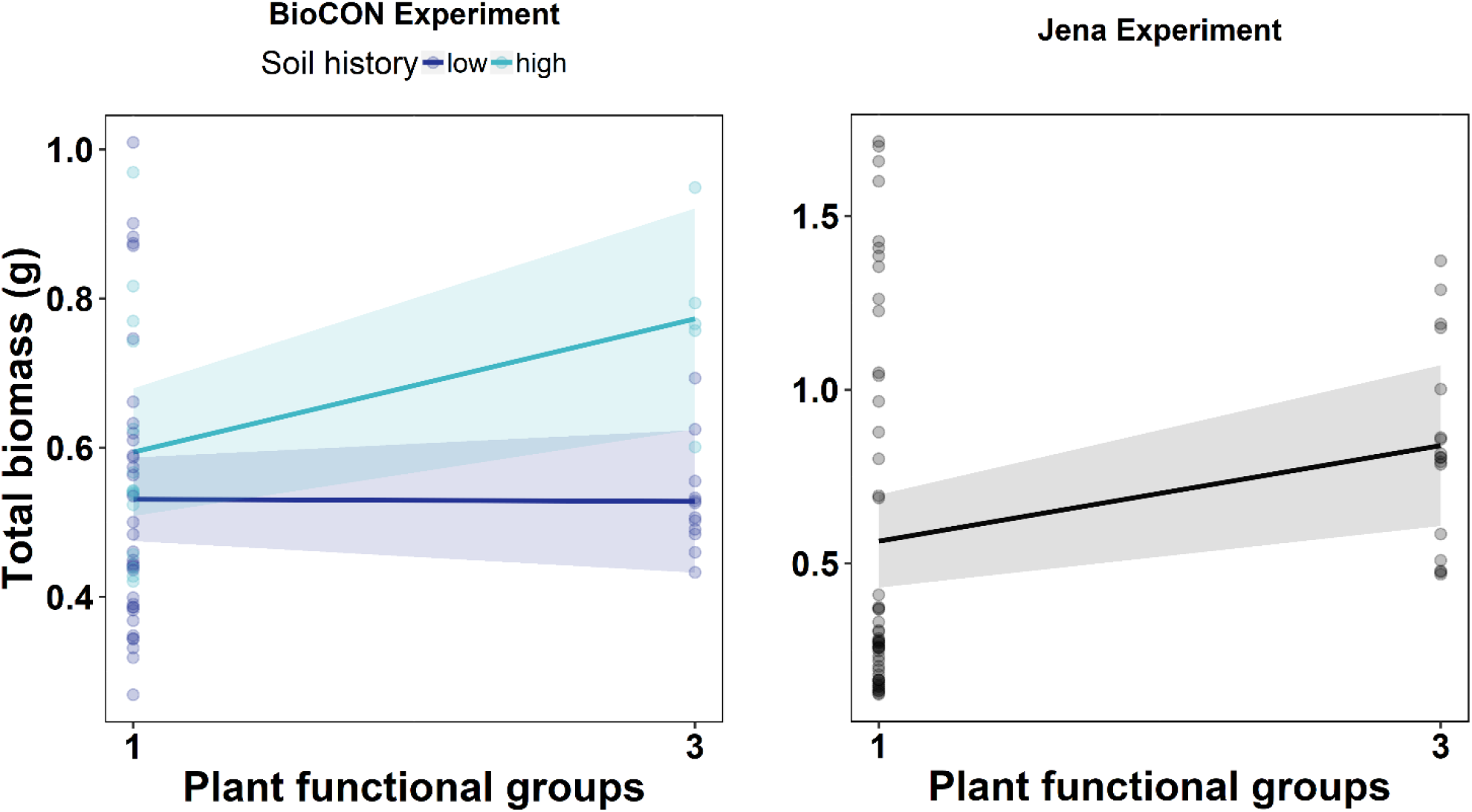
Total plant biomass in the microcosm experiment was explained by the interaction between soil history, i.e. soil conditioned by plant communities with either low- or high-diversity, and plant diversity in the microcosm experiment in the BioCON Experiment (marginally significant interaction effect, F_1,48_ = 3.179, p-value <0.1) and by plant diversity in the microcosm experiment in the Jena Experiment (F_1,49_ = 4.235, p-value < 0.05). Lines are linear mixed-effect model fits (bands indicate 95% confidence intervals). Points are microcosm level values.

### Effects of soil history on total plant biomass through differential mechanistic pathways

PSFs played a key role in determining total plant biomass in the microcosm experiment in the BioCON Experiment (Table S2, Fig. 3). Total biomass of plant species mixtures was higher in soil conditioned by high-than by low-diversity plant communities (Table S2, Fig. 3). Two mechanistic pathways were responsible for positive PSFs: nematodes richness and soil available N. Both nematode richness and soil available N were higher in soils conditioned by high-diversity than by low-diversity plant communities, with higher nematodes richness and available N increasing total plant biomass in plant mixtures in microcosms (standardized effects *via* nematode richness = 0.28 and *via* available N = 0.20). Positive PSFs in the BioCON Experiment were not fully explained by the measured mechanistic pathways, with a remaining significant path between soil history and total plant biomass in mixtures (standardized effect = 0.23).

**Figure 3.**
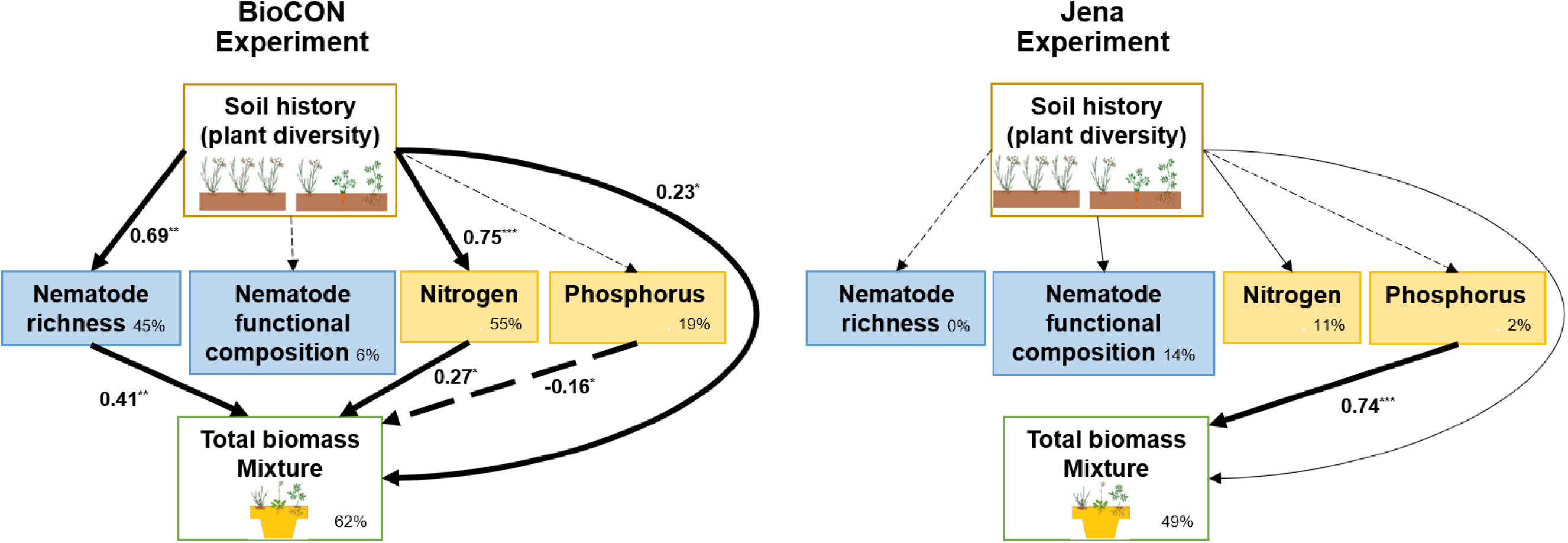
Structural equation models exploring the influence of soil history, i.e. soil conditioned by plant communities with either low- or high-diversity, on total biomass of plant species mixtures in the feedback experiment. Soils from the BioCON and Jena Experiments were used. Soil history may influence total biomass through biotic and abiotic mechanistic pathways, i.e. *via* estimated nematode richness, nematode functional composition, and available nitrogen and phosphorus concentrations; and/or through unmeasured mechanisms (direct path between soil history and total biomass in microcosms). The location of the plots where the soil was sampled in each biodiversity experiment was also included as a random effect. Nematode functional composition is based on a principal component analysis (PC2); PC2 represents a gradient from plant antagonists (plant-feeders) to plant growth facilitators (fungal-feeders) in both biodiversity experiments. In the BioCON Experiment, PC2 explained 22% of the variation, with positive values associated with a higher abundance of fungal-feeders and a lower abundance of plant-feeders. In the Jena Experiment, PC2 explained 34% of the variation, with positive values associated to a higher abundance of plant-feeders and omnivore nematodes and lower abundance of fungal feeder nematodes. Wide lines show significant effects (p-values <0.05*, <0.01**, or <0.001***), with solid and dotted black lines showing positive and negative effects, respectively. Explained variation (R^2^) is included for each response variable. Standardized path estimates are shown. The BioCON Experiment: Fisher’s C = 1.62, df = 2, p-value = 0.446, K = 23, n = 17 and the Jena Experiment: Fisher’s C = 7.35, df = 6, p-value = 0.29, K = 21, n = 17.

Soil available P affected total biomass in plant mixtures in the microcosm experiment in both, the BioCON and Jena Experiments (Tables S2 and S3, Fig. 2). Yet, the effect of available P (standardized effect in the BioCON Experiment = −0.16 and in the Jena Experiment = 0.74) was not related to PSFs associated to plant diversity in the conditioning phase. In the Jena Experiment, we found no evidence of PSFs on plant biomass in mixtures (Table S3, Fig. 3).

In the BioCON Experiment, PSFs did not influence total plant biomass of grasses growing alone in the microcosm experiment (Table S4, Fig. 4). In contrast, PSFs influenced the total plant biomass of forbs growing alone (standardized effect = 0.56). Specifically, total biomass of forbs increased in soil conditioned by high-diversity plant communities, suggesting negative PSFs in low-diversity communities. However, neither of the mechanistic pathways explained this negative PSF. Further, PSFs influenced total plant biomass of legumes growing alone *via* available N (standardized effect = 0.74). In other words, soil conditioned by high-diversity communities had higher available N than soil conditioned by low-diversity communities, with this difference in available N affecting total plant biomass of legumes (legume biomass increased with increasing concentrations of available N). Moreover, total biomass of legumes was negatively affected by soil history (standardized effect = −0.67).

**Figure 4.**
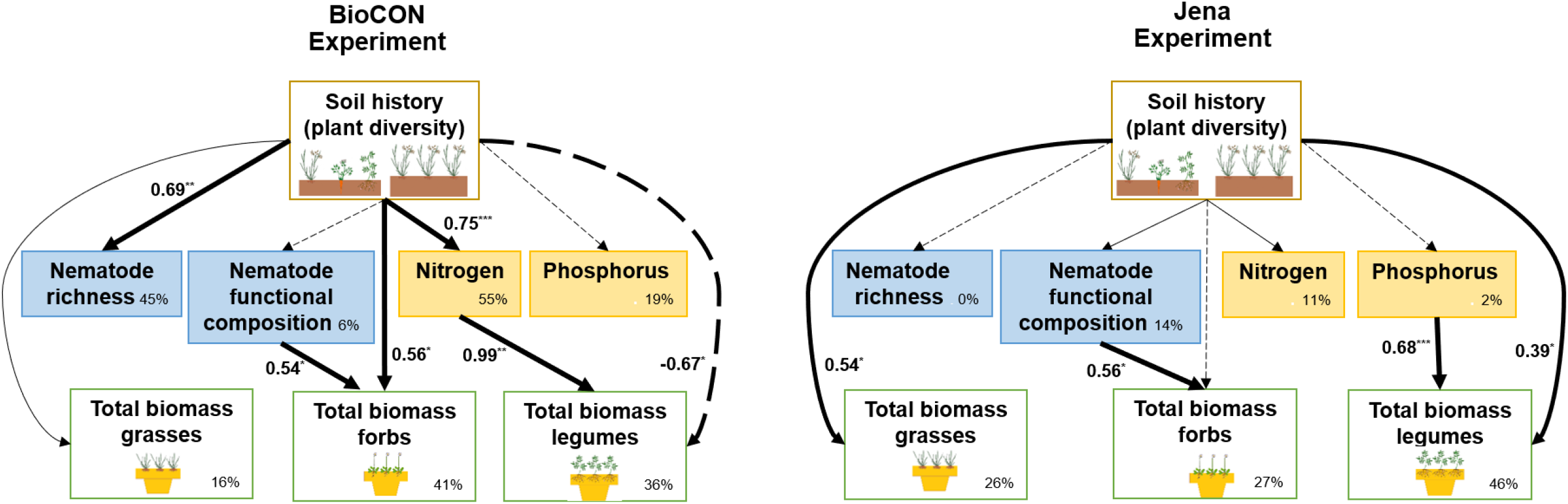
Structural equation models exploring the influence of soil history, i.e. soil conditioned by plant communities with either low- or high-diversity, on total biomass of plant communities containing only one functional group in the feedback experiment. Soils from the BioCON and Jena Experiments were used. Soil history may influence total biomass through biotic and abiotic mechanistic pathways, i.e. *via* estimated nematode richness, nematode functional composition, and available nitrogen and phosphorus concentrations; and/or through unmeasured mechanisms (direct path between soil history and total biomass in microcosms). The location of the plots where the soil was sampled in each biodiversity experiment was also included as a random effect. Nematode functional composition is based on a principal component analysis (PC2); PC2 represents a gradient from plant antagonists (plant-feeders) to plant growth facilitators (fungal-feeders) in both biodiversity experiments. In the BioCON Experiment, PC2 explained 22% of the variation, with positive values associated with a higher abundance of fungal-feeders and a lower abundance of plant-feeders. In the Jena Experiment, PC2 explained 34% of the variation, with positive values associated to a higher abundance of plant-feeders and omnivore nematodes and lower abundance of fungal feeder nematodes. Wide lines show significant effects (p-values <0.05*, <0.01**, or <0.001***), with solid and dotted black lines showing positive and negative effects, respectively. Explained variation (R^2^) is included for each response variable. Standardized path estimates are shown. The BioCON Experiment: Fisher’s C = 25.47, df = 20, p-value = 0.184, K = 30, n = 16 and Jena Experiment: Fisher’s C = 16.73, df = 20, p-value = 0.671, K = 30, n = 17.

In the Jena Experiment, PSFs influenced total biomass of grasses growing alone in the microcosm experiment (standardized effect = 0.54). In other words, grasses grew better in soils conditioned by high-than by low-diversity plant communities, suggesting negative PSFs in low-diversity communities. However, neither of the measured mechanistic pathways explained the PSFs on total biomass of grasses (Table S5, Fig. 4). While PSFs did not influence total biomass of forbs, PSFs influenced total biomass of legumes growing alone (standardized effect = 0.39).

Nematode functional composition affected total biomass of forbs growing alone in both, the BioCON and Jena Experiments (Tables S4 and S5, Fig. 4). Yet, their influence was not related to PSFs induced by plant diversity in the conditioning phase. In the BioCON Experiment, soils with a higher abundance of fungal-feeder and lower abundance of plant-feeder nematodes increased the total biomass of forbs (standardized effect = 0.54). In the Jena Experiment, soils with higher abundance of plant-feeders and omnivores as well as a lower abundance of fungal-feeder nematodes increased the total biomass of forbs (standardized effect = 0.56). In the Jena Experiment, available P affected total biomass of legumes growing alone, but this effect was not related to PSFs induced by plant diversity in the conditioning phase (standardized effect = 0.68).

### Effects of soil history on plant shoot and root biomass

The effects of PSFs on plant biomass varied across plant organs. In the BioCON Experiment, shoot biomass of plant species mixtures in microcosms was higher in soil conditioned by high-than by low-diversity plant communities, suggesting a positive PSF in more diverse communities (Tables S6, Fig. S3). This positive PSF acted *via* nematode richness (standardized effect = 0.34). Therefore, shoot biomass in mixtures was higher in soil with higher nematode richness. The effect of PSFs on shoot biomass of mixtures was not fully explained by the measured mechanistic pathways, which is why a significant path between soil history and shoot biomass of mixtures remained significant in the model (standardized effect = 0.52). PSFs also affected root biomass of mixtures (standardized effect = 0.65). However, PSFs on root biomass of mixtures were not related to neither of the measured mechanistic pathways.

In the Jena Experiment, shoot biomass of plant species mixtures in microcosms was higher in soil conditioned by high-than by low-diversity plant communities (standardized effect = 0.38, Tables S7, Fig. S3), which was in line with the findings in the BioCON Experiment. In contrast, PSFs associated to plant diversity did not influence root biomass of plant species mixtures in microcosms. Available P affected both, shoot and root biomass in mixtures (standardized effect for shoots = 0.65 and for roots = 0.79, Table S7, Fig. S4). However, the effect of available P was not related to PSFs associated to plant diversity.

In the BioCON Experiment, PSFs affected shoot biomass but not root biomass of grasses growing alone. Thus, shoot biomass was higher when grasses grew in soil conditioned by more diverse communities (standardized effect = 0.49, Table S8, Fig. S4). The effects of PSFs on shoot biomass of grasses were not explained by neither of the explored mechanistic pathways. The inverse pattern of that for grasses was observed for shoot and root biomass of forbs growing alone (Table S9, Fig. S4). While shoot biomass of forbs was not affected by PSFs associated with plant diversity, root biomass of forbs was. PSFs on root biomass acted *via* available N (standardized effect = 0.62). Therefore, root biomass was higher in forbs growing in soil conditioned by high-than by low-diversity plant communities, suggesting a negative PSF of low-diversity communities. PSFs affected both shoot and root biomass of legumes growing alone *via* available N (standardized effect = 0.71 for shoots and 0.74 for roots; Table S10, Fig. S4). For root biomass of legumes, PSFs were not fully explained by the mechanistic pathways (standardized effect = -0.78).

In the Jena Experiment, PSFs associated to plant diversity did not affect shoot biomass but affected root biomass of grasses growing alone (standardized effect for root biomass = 0.52, Table S11, Fig. S4). However, the influence of PSFs on root biomass was not explained by neither of the measured mechanistic pathways. PSFs associated to plant diversity affect neither shoot nor root biomass of forbs (Table S12, Fig. S4). For legumes growing alone, while PSFs did not influence shoot biomass, root biomass increased in soils conditioned by more diverse plant communities (standardized effect = 0.41, Table S13, Fig. S4).

In the Jena Experiment, shoot biomass (of forbs and legumes) and root biomass (of legumes) were affected by biotic and abiotic conditions. Yet, these effects were not linked to PSFs associated to plant diversity. Specifically, shoot biomass of forbs increased in soils with higher abundance of plant feeding nematodes and omnivores and lower abundance of fungal-feeders (standardized effects = 0.64). Shoot and root biomass of legumes increased with available P (standardized effect = 0.71 for shoots and 0.56 for roots).

### Effects of soil history on plant biomass: the role of plant functional identity

In the BioCON Experiment, negative values of the PSF index indicated that plant communities with a single plant functional group grew significantly worse (for total and root biomass) in soil conditioned by low-diversity communities with the same plant functional group as compared to low-diversity communities conditioned by other plant functional groups (Fig. 5). In the Jena Experiment, a similar pattern was observed, but it was not statistically significant.

**Figure 5.**
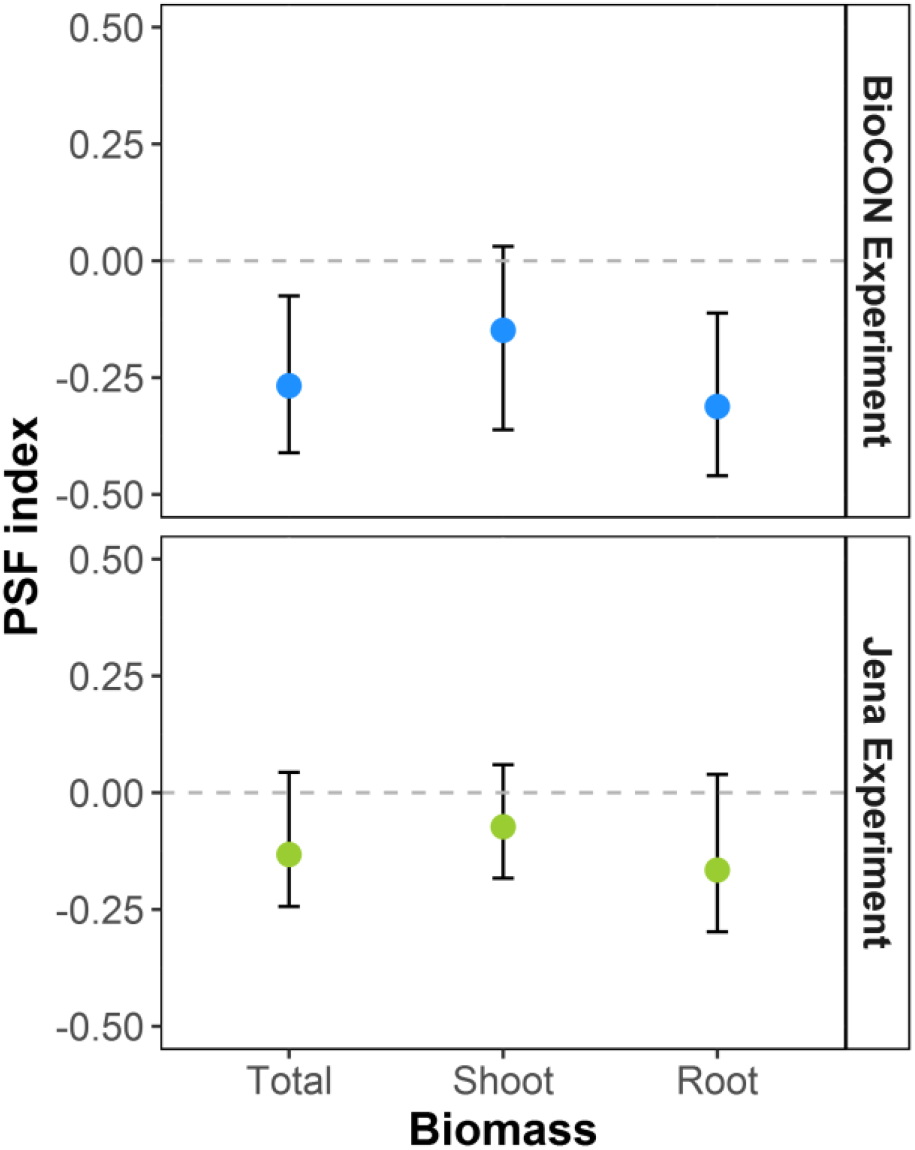
Effects of soil conditioned by the same functional group *vs*. soil conditioned by another plant functional group on total, shoot, and root biomass of low-diversity communities in the BioCON and in Jena Experiments. PSF index (mean ± 95% CI) is defined as the biomass of a low-diversity plant community matching in their plant functional composition between the field (conditioning phase) and the microcosm experiment (BiomassM^1^) (feedback phase) minus the mean of the biomass of low-diversity communities that did not have the same plant functional composition in the field as in the microcosm experiment (BiomassM^2^), divided by Biomass M^2^. Negative values indicate that plant communities with a single plant functional group grew worse in soil conditioned by low-diversity communities with the same plant functional groups, compared with low-diversity communities conditioned by other plant functional groups.

### Effects of soil history on plant diversity effects on plant biomass

Soil history changed not only plant biomass *per se* but also influenced plant diversity effects in the feedback phase in the BioCON Experiment (Fig. 6). Particularly, positive plant diversity effects on total and shoot biomass were observed when plant communities grew in soil conditioned by high-but not by low-diversity plant communities. In the Jena Experiment, positive plant diversity effects in the feedback phase on total, shoot, and root biomass were observed independent of the soil history.

**Figure 6.**
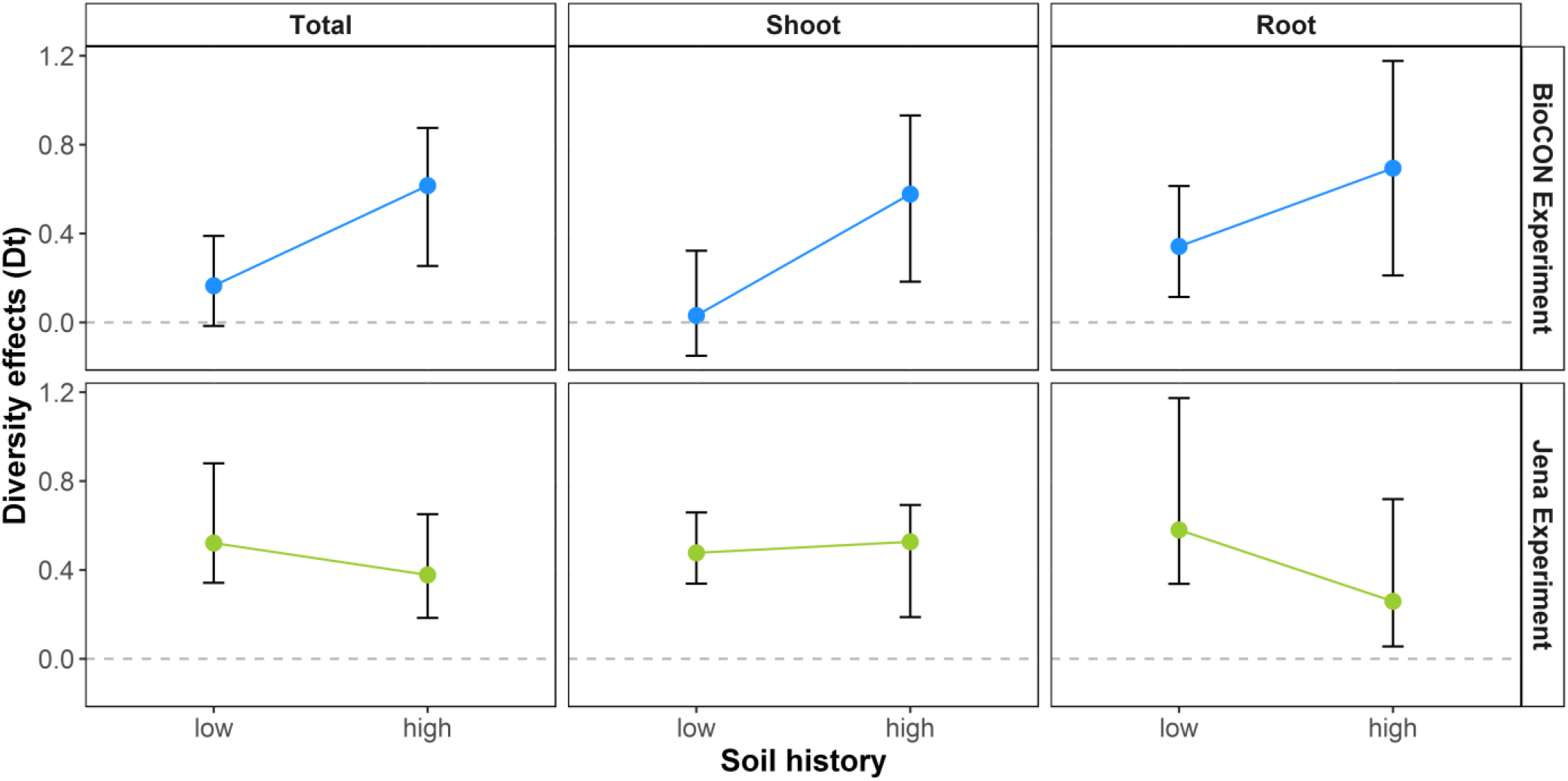
Soil history, i.e. soil conditioned by plant communities with either low or high plant species diversity, effects on plant diversity effects on total, shoot, and root biomass in the microcosm experiment (mean ± 95% CI) in the BioCON and Jena Experiments. Non-additive effects are considered to occur when the 95% CI do not overlap with zero.

## Discussion

This is one of the first studies that disentangles potential mechanistic pathways underlying diversity-dependent PSFs on primary productivity and plant diversity effects in long-term biodiversity experiments. Our results reveal that PSFs may drive positive plant diversity effects on ecosystem functioning by both positive PSFs of soil conditioned by high-diversity plant communities and negative PSFs of soil conditioned by low-diversity plant communities.

On the one hand, PSFs of soil conditioned by high-diversity plant communities in the BioCON Experiment mainly influenced diverse plant communities and not communities with a single plant functional group in the feedback phase. Therefore, PSFs in the BioCON Experiment not only affected plant productivity but also explained positive plant diversity effects in our microcosm experiment. On the other hand, low-diversity communities were negatively affected by growing on soil conditioned by their own plant functional group, especially in the BioCON Experiment, suggesting that negative effects associated with low-diversity communities may be amplified when plant species are functionally similar. In contrast, plant-plant interactions, and not plant-soil interactions, drove diversity effects on primary productivity in the Jena Experiment. Taken together, these results suggest that primary productivity and plant diversity effects on this ecosystem function can be driven by plant-soil and plant-plant interactions, with the main driver differing between experimental contexts.

Positive PSFs on plant productivity emerge due the influence of plant diversity of both the community conditioning the soil and the community growing in the conditioned soil. The absence of positive PSFs of high-diversity plant communities in other studies (Yang *et al*. 2015) may be the result of assessing PSFs at the species level and not at the community level, such as done in the present study. High-diversity communities may positively affect plant productivity by diversifying the abiotic and biotic biotope characteristics that facilitate plant growth (Hooper *et al*. 2000; Eisenhauer *et al*. 2012a). Yet potential mechanisms required to capitalize on this positive PSF may be present only in high-diversity communities and not in low-diversity communities (Loreau & Hector 2001, Tilman et al. 2014).

The mechanisms underlying diversity-dependent PSFs on primary productivity are likely interconnected and dynamic, increasing the challenge of disentangling confounding factors in order to establish causality. For instance, plant diversity of the soil conditioning community increased productivity of forbs and legumes *via* available N. Although it is not surprising that an increase of available N leads to an increase in plant productivity (Craven *et al*. 2016), lower N availability was associated with soils previously conditioned by forbs and legumes (Fig. S5). In other words, biomass of forbs and legumes was lower in soil conditioned by species belonging to their own plant functional groups. Thus, negative PSFs of forbs and legumes likely caused the observed plant identity effects on PSFs in our microcosm experiment for the BioCON Experiment. Interestingly, the same pattern has been observed in the field, in which a decrease in root biomass in forbs and legumes was suggested as a potential mechanism explaining a temporal decrease in available N (Mueller *et al*. 2013). Although nutrient depletion (in low-diversity communities) and/or enrichment (in high-diversity communities) may explain the observed PSFs, other potential mechanisms are also possible. For instance, pathogens could exacerbate the decrease of root biomass (and subsequent nutrient depletion), thereby reducing either the recruitment or the performance of specific plant species or functional groups (Fornara & Tilman 2008; Petermann *et al*. 2008; Reich *et al*. 2012).

Inconsistent PSFs on primary productivity between biodiversity experiments may reflect differences in the environmental context, such as soil fertility. Positive effects of plant species diversity on available N may be more pronounced in soils with low fertility, such as the sandy and organic-poor soil in the BioCON Experiment. Positive effects of plant species diversity on NH4 may emerge because of an increase over time of soil carbon concentration in more diverse communities. Subsequently, cation exchange sites in the soil organic matter may be filled by NH_4_ ions, resulting in an increase of NH_4_ in more diverse communities (Fornara & Tilman 2008; Mueller *et al*. 2013). Temporal dynamics of nutrients can be also observed in more fertile soils as in the Jena Experiment (Oelmann *et al*. 2011). However, weaker effects may emerge due to either constraints in the potential of plant species diversity to significantly increase nutrient availability in fertile soils or the absence of specific mechanisms driving the contrasting effects of high- and low-diversity plant communities on available nutrients, e.g., differences in root biomass across plant richness levels (Bessler *et al*. 2009). It is also possible that PSFs in more fertile soils may not affect plant productivity *per se* but rather the recruitment of specific species (LaManna *et al*. 2016). Also changes in primary productivity in fertile soils may be mainly driven by other resources (Hautier *et al*. 2009) or by plant community evolutionary changes *via* selection (van Moorsel *et al*. 2018).

Dissimilarity between drivers of plant diversity effects on ecosystem functioning between biodiversity experiments may emerge as biotic interactions likely co-vary with environmental conditions. Biotic interactions can be impacted by soil chemical and physical characteristics that directly influence soil communities (Hassink *et al*. 1993; Wardle *et al*. 2004) and/or modify plant-microbial competition and facilitation (Kuzyakov & Xu 2013; Johnson & Rasmann 2015). The fertility of the ecosystem may shape plant community composition as well as impact brown and green energy channels, with these changes cascading to ecosystem processes (Wardle *et al*. 2004). Positive effects of plant growth facilitators are often more pronounced in nutrient-poor than in nutrient-rich soils (Johnson 2009; van Groenigen *et al*. 2014). Differences in soil characteristics like soil texture affect the grazing activity of nematodes (bacterial-feeders), resulting in higher soil N mineralization rates in sandy than in clay soils (Hassink *et al*. 1993). Further, effects of plant antagonists may differ between soil fertility levels through changes in compensatory plant growth responses to herbivory. A reduction of compensatory effects in nutrient-poor soils may amplify the negative effects of herbivory, affecting not only plant growth but also herbivore-plant-microbial interactions (Bardgett & Wardle 2003). Thus, soil fertility may co-determine plant-soil biotic interactions as well as the consequences for ecosystem functioning.

In conclusion, temporal changes in the B-EF relationship may emerge because of diversity-dependent PSFs. However diversity-dependent PSFs rely on the experimental context, reinforcing the notion that environmental conditions are likely to play a key role in determining not only the patterns (Guerrero-Ramírez *et al*. 2017), but also the mechanisms underlying temporal changes in the B-EF functioning relationship in grasslands. Since plant species diversity can modify ecosystem functioning through both plant-plant interactions as well as through soil legacy effects (Zuppinger-Dingley *et al*. 2016), negative effects of plant diversity loss on ecosystem functioning may persist or even increase over time.

## Acknowledgements

Special thanks to Alfred Lochner for his support during all the experimental phase and Petra Hoffmann (UFZ/iDiv, Physiological Diversity) for her support during the chemical analyses. We thank Kally Worm, Anne Ebeling, Simone Cesarz, Silke Schroeckh, and Katja Steinauer for helping us make this experiment possible and the gardeners and coordinators of both long-term biodiversity experiments for their work to maintain the plant diversity gradient. We thank Dylan Craven for his help with the statistical analyses and providing comments on a previous version of this manuscript. We thank Andrew Barnes and Jon Lefcheck for their input on structural equation models. This work was funded by the German Research Foundation (Deutsche Forschungsgemeinschaft, DFG) in the frame of the Emmy Noether research group (Ei 862/2) to NE. Further support came from the German Centre for Integrative Biodiversity Research (iDiv) Halle-Jena-Leipzig, funded by the DFG (FZT 118). The Jena Experiment is financed by the Deutsche Forschungsgemeinschaft (FOR 1451). Support for the BioCON Experiment came from the US National Science Foundation (NSF) Long-Term Ecological Research (DEB-9411972, DEB-0080382, DEB-0620652 and DEB-1234162), Biocomplexity Coupled Biogeochemical Cycles (DEB-0322057), Long-Term Research in Environmental Biology (DEB-0716587, DEB-1242531) and Ecosystem Sciences (NSF DEB-1120064) Programs; as well as the US Department of Energy Programs for Ecosystem Research (vDE-FG02-96ER62291) and National Institute for Climatic Change Research (DE-FC02-06ER64158).

